# Efficient exact associative structure for sequencing data

**DOI:** 10.1101/546309

**Authors:** Camille Marchet, Mael Kerbiriou, Antoine Limasset

## Abstract

**Motivation:** A plethora of methods and applications share the fundamental need to associate information to words for high throughput sequence analysis. Doing so for billions of *k*-mers is commonly a scalability problem, as exact associative indexes can be memory expensive. Recent works take advantage of overlaps between *k*-mers to leverage this challenge. Yet existing data structures are either unable to associate information to *k*-mers or are not lightweight enough.

**Results:** We present BLight, a static and exact data structure able to associate unique identifiers to *k*-mers and determine their membership in a set without false positive, that scales to huge *k*-mer sets with a low memory cost. This index combines an extremely compact representation along with very fast queries. Besides, its construction is efficient and needs no additional memory. Our implementation achieves to index the *k*-mers from the human genome using 8GB of RAM (23 bits per *k*-mer) within 10 minutes and the *k*-mers from the large axolotl genome using 63 GB of memory (27 bits per *k*-mer) within 76 minutes. Furthermore, while being memory efficient, the index provides a very high throughput: 1.4 million queries per second on a single CPU or 16.1 million using 12 cores. Finally, we also present how BLight can practically represent metagenomic and transcriptomic sequencing data to highlight its wide applicative range.

**Availability:** We wrote the BLight index as an open source C++ library under the AGPL3 license available at github.com/Malfoy/BLight. It is designed as a user-friendly library and comes along with code usage samples.

## 1 Introduction

Tremendous, ever-growing amounts of DNA and RNA reads are made available through high-throughput sequencing methods. Single RNA, DNA, or metagenome and metatranscriptome samples can contain up to billion reads each and the NIH Sequence Read Archive (SRA) [1] now gathers peta-bases of sequences. Working on such collections of samples is a fundamental challenge for associative indexation schemes that label each sequence of interest, e.g., all fixed-length words (*k*-mers), with a unique identifier. This fundamental block is a corner stone for a large spectrum of methods in bioinformatics: genome assembly [2], efficient overlap detection among large sequences [3], quick quantification of transcriptomes [4], sequence search in large sequences collections [5], variant detection [6]; and can be identified as a generic need in large scale sequence analysis. Even after the assembly step, indexing very large genomes (e.g., *Pinus taeda* [7] with 20 Gbp or *Ambystoma mexicanum* [8] with 32 Gbp) or metagenomes remain a serious difficulty. Numerous efforts focus on designing data structures performing the generic task of associating information to words from studied sequences that can scale to these magnitudes. The main difficulty remains to design data structures that can handle billions of distinct *k*-mers so that they can scale up to most voluminous instances such as genomes, genome collections, or metagenomics datasets. There are three strategies commonly considered. The first one is to index fixed-size words (*k*-mers) from sequences in structures that enable presence/absence queries. This strategy often relies on Bloom filters [9] although such data structures can only determine the membership of arbitrary *k*-mers and cannot associate information to them. Recently Bloom filters, or other probabilistic sets, have been used to search sequences in thousands of indexed raw datasets [10], or for assembly [11, 12]. A second strategy uses full-text indexes to localize words of arbitrary length in a set of sequences. They commonly rely on FM-indexes [13]. These methods offer extremely memory-efficient indexes but at a high construction cost and a reduced throughput compared to hash-based methods. Finally, some data structures propose general associative indexes. Based on hash tables [6] and/or filters [5], they allow to store (*k-mer, value*) pairs. This way, *k*-mers can be associated with pieces of information of any nature, for instance, with their original dataset(s) [4], or counts [14]. The presented work pertains to this latter category. Pioneer works based on associative structures such as Cortex [6] illustrate the high hash tables cost that can require more than a dozen bytes per *k*-mer. Cortex enhances the de Bruijn graph built from the union of different strains, species or samples, by associating their datasets of origin to each *k*-mer of the graph using a hash table. However, Cortex hash scheme was designed mainly for speed and cannot scale up to more than a dozen datasets. Such difficulty motivated recent improvements [15, 16], either based on practical and efficient implementation of a minimal perfect hash function (MPHF) [17] or FM-index.

Indexes build upon MPHFs present fast queries using less than 4 bits per *k*-mer for the MPHF itself. However, MPHFs are bijective functions between a key set and its value set, but do not represent sets. Additional information is necessary to reject keys absent from the indexed set (alien keys). In a previous work, we proposed adding a structure to associate to each *k*-mer a hash-based fingerprint [18], to obtain a probabilistic associative index. The structure has some false positives due to hash collisions, which depend on the fingerprint size. More recently, a method [16] took advantage of the possibility to assemble *k*-mers using *compacted de Bruijn graphs* [19, 20] in the tool Pufferfish. They store the position of each *k*-mer in the set of assembled sequences and can reject alien *k*-mers by seeking their sequence at the position indicated by the index.

Representing *k*-mers sets benefits from recent works that are valuable to indexation tasks. Recently, spectrum-preserving string sets (SPSS) [21] were defined as an exact representation of a multiset of *k*-mers coming from a set of strings of length ≥ *k*. In the literature, SPSS narrow this definition to the representation of a *k*-mer set and forget the *k*-mer multiplicities. According to this definition and as noticed in Pufferfish, de Bruijn graphs are relevant SPSS since they collapse redundancy in their vertices representing the *k*-mers set. In this work, we will explore different SPSSs for indexing *k*-mers. Moreover, there exist numerous efficient de Bruijn graph representations (succinct data structures such as BOSS [22], efficient representations of de Bruijn graphs vertices such as DBGFM [23] or deGSM [24]). However, they differ from this work’s scope since they are not associative structures, or at least not implemented as such.

In this contribution, we propose a novel, exact associative structure for *k*-mers dubbed BLight, able to scale to large datasets while being particularly memory efficient and fast. Although it uses an MPHF, this structure is deterministic and yields no false positives at the query. It enabled the indexation of the 18 billion of 31-mers from the axolotl genome using 62.4GB of RAM (≈ 27 bits per *k*-mer) in 76 minutes. The BLight index performed 1.4 million queries per second on a single core and more than 16 million queries per second using 12 cores on the human genome. Contrary to works dedicated to a particular application (colored de Bruijn graphs [5], quantification [4]), the BLight index is a generic associative index that can fit a wide range of different purposes that can associate any information to *k*-mers. To this extent, we designed it as a user-friendly library with sample code usage to facilitate its integration to various projects. We demonstrate its performances on different datasets to illustrate its potential applications on various issues.

## 2 Methods

### 2.1 Preliminary definitions

For simplicity’s sake we describe the methods without taking into accounts reverse-complement, in practice a *k*-mer and its reverse-complement will always be considered identical.

#### Definition 1.

***de Bruijn graph:*** *The de Bruijn graph is a directed graph G*_*k*_ =(*V,E*) *where V is a set of k-mers. For u,v ∈ V*, (*u,v*) ∈ *E if and only if the suffix of size k* − 1 *of u is equal to the prefix of size k* − 1 *of v. Note that this definition is node-centric: the k-mer set is equivalent to the node set and the set of edges can be directly inferred from the nodes set*.

#### Definition 2.

***Unitig:*** *Given a de Bruijn graph G*_*k*_, *a unipath is a maximal-length linear path (sequence of distinct nodes) s* =[*u*_0_,…,*u*_*i*_,…,*u*_*n*_] *such that*

- *for each* 0 ≤ *i* ≤ *n* − 1, *the edge e* =(*u*_*i*_,*u*_*i*+1_) ∈ *E;*
- *in and out-degrees are equal to 1 for each u*_*j*_ *such that* 1 ≤ *j* ≤ *n* − 1;
- *if n>*1 *the out-degree of u*_0_ *is 1;*
- *if n>*1 *the in-degree of u*_*n*_ *is 1*.

*Given a unipath* [*u*_0_,…,*u*_*i*_,…,*u*_*n*_], *a unitig corresponds to the concatenation of nodes in the unipath, such that u*_0_*’s string is concatenated with each last character of u*_*i*_ *in order, for* 1 ≤ *i* ≤ *n. Thus, the unitig is a string of length k*+*n. Disconnected k-mers of G*_*k*_ *are also unitigs*.

#### Definition 3.

***Compacted de Bruijn graph:*** *The directed graph where nodes are unitigs, and edges are k*−1*-overlaps between two nodes sequences, is called a compacted de Bruijn graph [20]*

#### Definition 4.

***Minimizer:*** *Minimizers were defined in [25]. The m-minimizer of a sequence S is the smallest substring of size m in S, according to some order. In this work, we compute m-minimizers from k-mers, and a hash function defines the order on m-mers*.

### 2.2 Outline

#### 2.2.1 Inputs/outputs

For now, we assume that we construct the BLight index from the unitigs of the compacted de Bruijn graph constructed from a *k*-mer set. They can be constructed efficiently with Bcalm2 [20] from any FASTA/FASTQ file compressed or not. Later we will see that we can perform the construction from other types of sequences. For each *k*-mer present in the input graph, the BLight index returns a unique identifier *i* ∈[1,*N*] with *N* the total number of *k*-mers, and −1 for any other *k*-mer.

#### 2.2.2 BLight index construction

First, we split the graph into several *k*-mer sets called buckets, according to the *k*-mer minimizers (See Figure 1). A bucket handles all *k*-mers getting a given minimizer, hence for a minimizer size *m* we construct 4^*m*^ buckets. At first, each bucket contain its *k*-mer sequences encoded in binary. Once filled, we build for each bucket a specific MPHF from its *k*-mer set. Thus, for a given bucket the MPHF returns a unique identifier for each of its *k*-mer. This way we can associate a unique identifier to each indexed *k*-mer. We also use this identifier to locate each *k*-mer in its associated bucket. To do so we associate to each *k*-mer its position in the bucket sequences. Thereby, during a query, we can check if the query *k*-mer is effectively in the index and not an alien *k*-mer. In total, a bucket contains its *k*-mer sequences (binary encoded), an MPHF associating to each *k*-mer its position, and the *k*-mer positions themselves. Each *k*-mer position needs 𝒪 (*log*_2_(*bucket size*)) bits to be encoded, the *k*-mer sequences need two bits per nucleotide and the MPHF cost roughly 4 bits per *k*-mer. We choose to encode both the positions and the *k*-mers using bit-vectors to optimize the required space.

**Figure 1.**
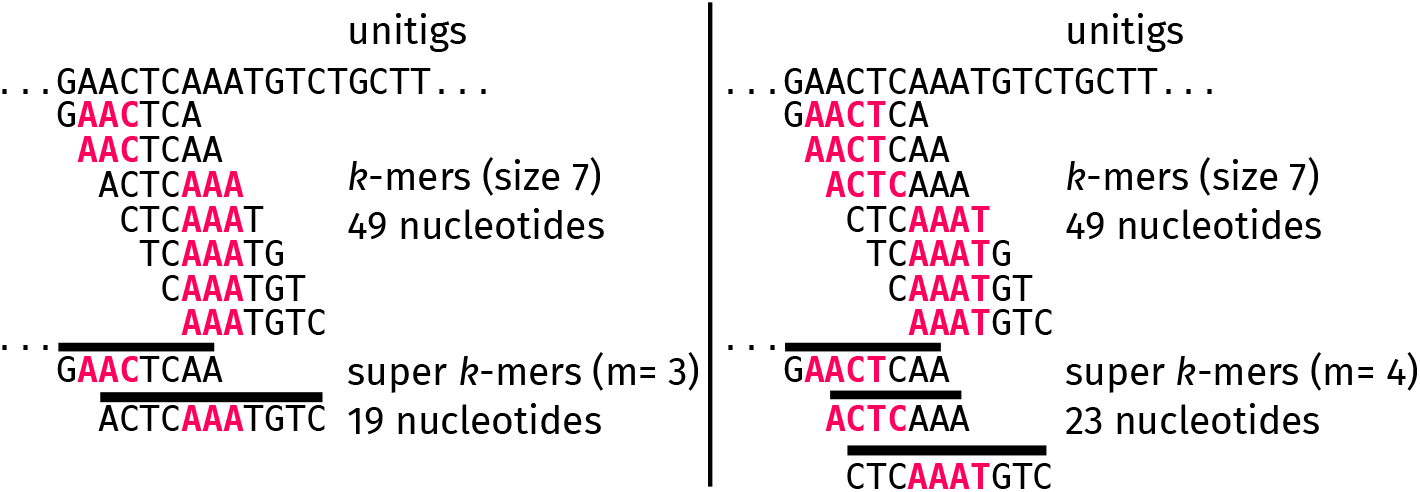
Example of splitting a unitig into super-*k*mers. The set of 7-mers uses 49 nucleotides to represent the same information as in the unitig. Left: with minimizers (highlighted) of size 3. The super-*k*-mers from unitigs use 19 nucleotides. Right: with minimizers of size 4. Less *k*-mers share the same minimizer, leading to a fragmentation of the super-*k*-mers, and more nucleotides to represent the same set.

#### 2.2.3 Query

The query has two steps (Figure 3): first, we select the relevant bucket according to the query *k*-mer minimizer. Second, we query the MPHF with the *k*-mer. The MPHF can return false positives at the query but no false negatives. For an existing *k*-mer, the MPHF returns its identifier that points to its position in its bucket (Figure 3 (a)). For an alien *k*-mer, the MPHF either returns no information, meaning the *k*-mer is not present, or it falsely returns a position (Figure 3 (b)). Then, when the MPHF points to a position *p*, we check it in the sequences bucket. This way, we check that the correct *k*-mer sequence is at the indicated position. If the *k*-mer sequence is not present or the MPHF returns no information, we return − 1 to indicate this *k*-mer is not in the index. Thus, it is essential to notice that, despite relying on the probabilistic MPHF response, our structure always returns exact answers.

**Figure 3.**
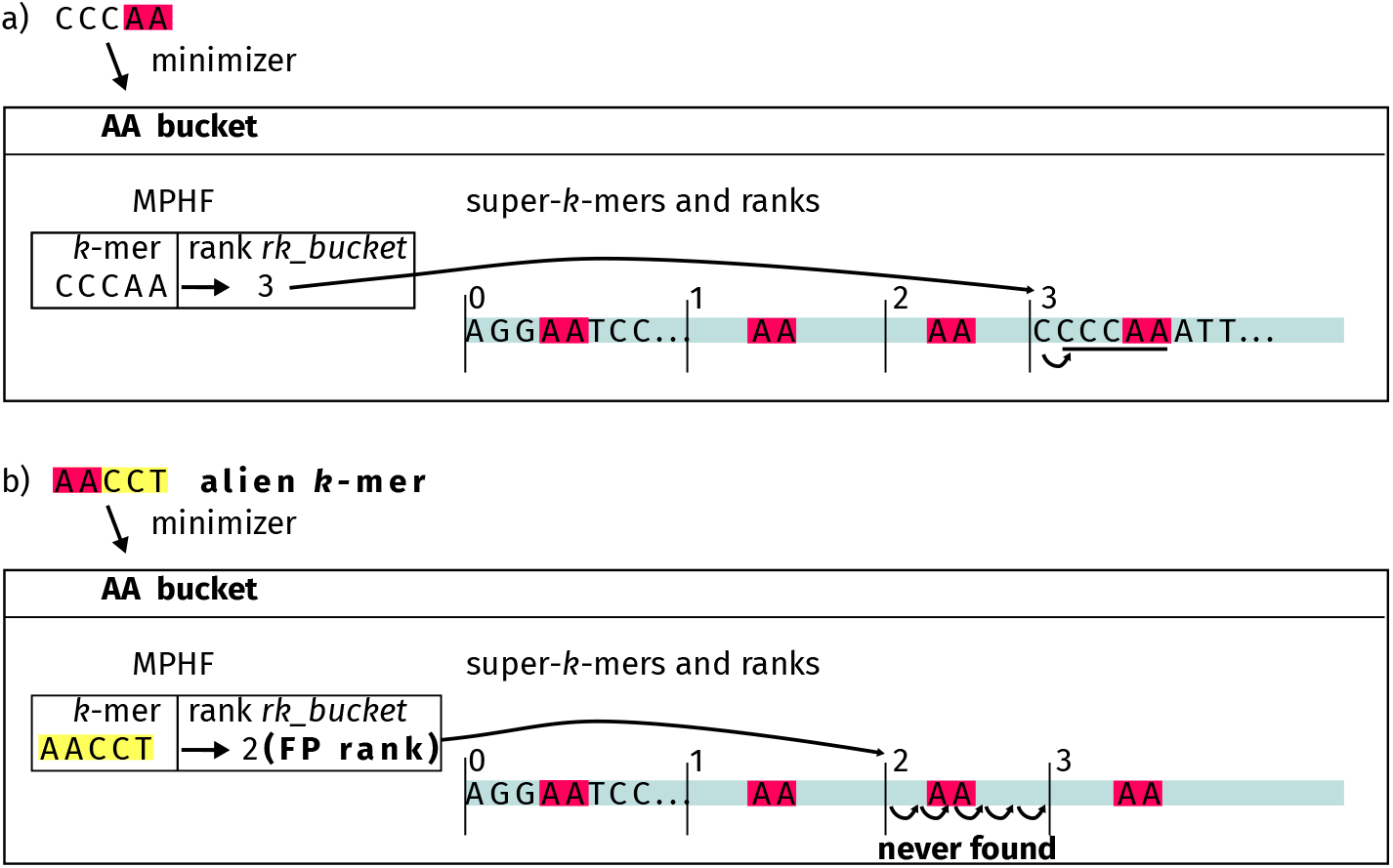
Queries in the index: (a) query of an indexed *k*-mer, (b) query of an alien *k*-mer. We represent minimizers in dark grey. Concatenated super-*k*-mers of a bucket are in light grey. In (a), after finding the appropriate bucket (box) using the *k*-mer’s minimizer, the *k*-mer’s super-*k*-mer rank is returned, and we search the *k*-mer sequence from that position (underlined). In (b), a false positive rank (FP) is returned for the alien *k*-mer; hence it cannot be found in the given super-*k*-mer.

### 2.3 Data structure description

#### 2.3.1 Super-*k*-mers

In order to represent *k*-mers from a bucket in a memory-efficient way, we rely on the notion of super-*k*-mers, that appeared in [26]. The (ordered) list of its *k*-mers can represent any sequence. In practice, we observe that consecutive *k*-mers often share the same minimizer. Therefore a list of *x* successive *k*-mers from a sequence sharing the same minimizer can be encoded as a word of length *k*+*x* − 1, called a super-*k*-mer (Figure 1). This technique permits an efficient partition of a multi-set of *k*-mers that use fewer sequences than raw *k*-mers. It is mainly used to compute *k*-mers abundances within a dataset [27].

In *k*-mer counters [26, 27], the super-*k*-mers are computed from reads. In this paper, in order to rely on the partitioning property, we extract super-*k*-mers from unitigs instead of reads (Figure 2). Since we have no duplicate *k*-mers in our unitigs, those super-*k*-mers represent a set of *k*-mers instead of a multi-set. We use super-*k*-mers in order to store *k*-mers more efficiently in each bucket. It is interesting to note that most tools define super-*k*-mers as maximal list of consecutive *k*-mers sharing a minimizer. For technical reasons, super-*k*-mers can also be non-maximal. Despite being a sub-optimal encoding, a very large *k*-mer list could be encoded in several super-*k*-mers, with no impact on the represented set, nor on the *k*-mers content.

**Figure 2.**
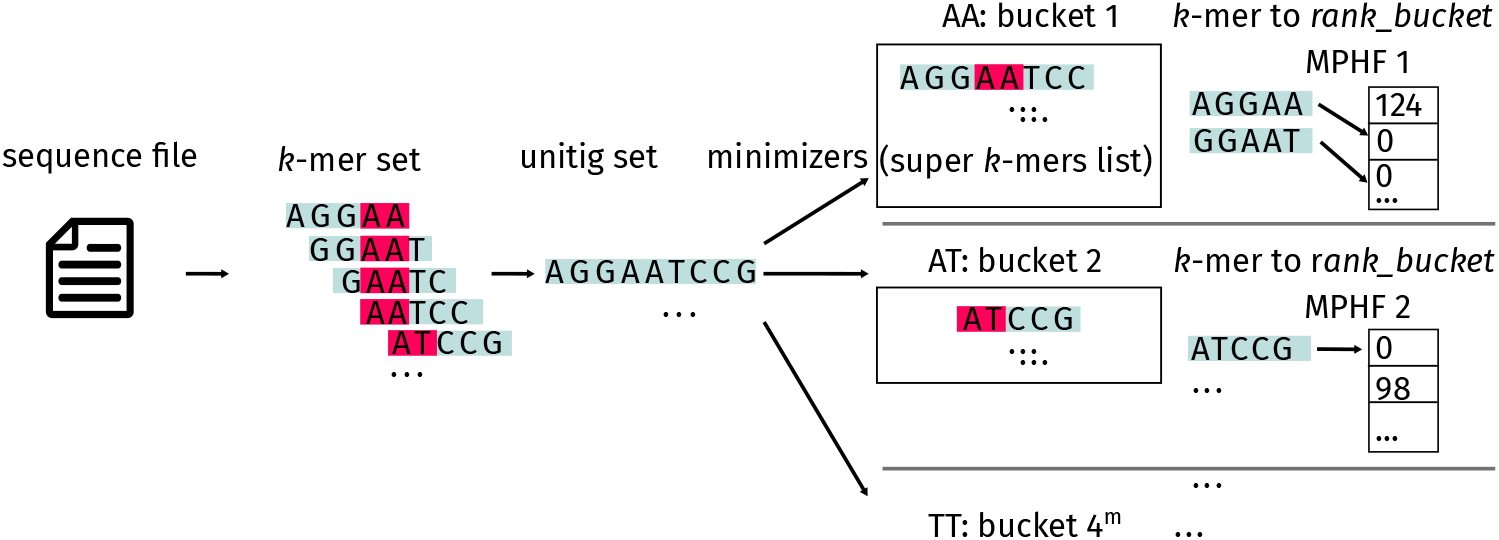
Overview of BLight’s method presented in Figure 2. We build a compacted de Bruijn graph, from the initial sequence file. We split unitigs from the graph into super-*k*-mers and put them in buckets, using minimizers (highlighted in grey). When querying, a *k*-mer is looked-up into the relevant bucket using its minimizer. Each bucket is indexed by an MPHF that associates a unique identifier to each *k*-mer. We also use this identifier to associate *k*-mers to their ranks in the bucket. This way, *k*-mers can be retrieved in the bucket during the query.

We write their sequences and their associated minimizers on the disk in gzipped FASTA format. This way, each bucket can be either processed in parallel to accelerate the index construction or independently to limit the memory used.

#### 2.3.2 Rank structure

In practice, instead of storing each *k*-mer position, we store the identifier of the *k*-mer’s super-*k*-mer. This solution allows using *log*_2_(*number superkmer*) instead of *log*_2_(*number nucleotides*) bits per *k*-mer in the index, which can represent a difference up to 6 bits per *k*-mer on our experiments. To recover the correct super-*k*-mers in constant time, we use a compressed rank structure^1^. We construct a bit-array for each bucket, associating a bit to each *k*-mer. A bit is set to 1 if the corresponding *k*-mer is the first of its super-*k*-mer. This bit array is compressed and indexed by the rank structure that allows the localization of a given super-*k*-mer in its bucket from the following formula: *position ith superkmer* =*select*(*i*)+*i* × (*k* − 1) as each super-*k*-mer is followed by *k* − 1 nucleotides that are not indexed *k*-mers. Once the corresponding super-*k*-mer is selected, we check among its *k*-mers if the query *k*-mer is effectively present. If not, it means that the query *k*-mer is an alien key. Even if the amount of *k*-mers forming a super-*k*-mer is unlikely to be above 2 ×*k* − *m*, we choose to bound it to ensure that queries take constant time in practice.

#### 2.3.3 Minimizer size and fragmentation

The number of buckets increases exponentially (4^*m*^) as the minimizer size *m* increases. As a consequence, buckets contain less *k*-mers (and super-*k*-mers) at higher *m* values, which means that the bit-vectors used to encode positions in each bucket are also globally reduced (as each *k*-mer position requires *log*_2_(*number superkmer*) bits to be encoded). Thus, increasing the minimizers’ size permits to encode *k*-mers positions in a more efficient way.

However, increasing the number of buckets also increases the structure weight, as each bucket requires some allocated memory independently of the input. As the number of buckets is exponential according to *m*, it can induce a massive memory consumption when *m* is above 10. Moreover, a higher *m* generate more, smaller super-*k*-mers [28] and this fragmentation raises the total amount of nucleotide necessary to represent the *k*-mer set (See Figure 1).

Another source of fragmentation is the graph complexity. Since super-*k*-mers can only be smaller than unitigs, a tangled graph containing many small unitigs will produce many small super-*k*-mers that will be costly to encode and index. Therefore the graph fragmentation can diminish the index efficiency by fragmenting unitigs. We point out that BLight can directly work on any SPSS formatted in FASTA format to mitigate this downside as we can build the super-*k*-mers from any set of sequences without duplicated *k*-mers. Working on sequences longer than unitigs is especially useful for large and repetitive genomes where the compacted de Bruijn graph is fragmented into a high amount of small unitigs.

#### 2.3.4 Hierarchical bucketing

In practice, a uniform partition is hard to obtain. The bucket size distribution is highly unbalanced [23]. The smallest minimizers create large buckets, while some buckets can be empty. The largest buckets are notably costly to index because all their *k*-mers will use 𝒪 (*log*_2_(*bucket size*)) bits. We propose a novel strategy called a hierarchical bucketing scheme to tackle this problem. Each bucket will contain smaller *k*-mer sets (sub-buckets) associated with several larger minimizers. By randomly combining smaller sub-buckets, we expect to obtain smaller largest buckets, as it is unlikely to group large sub-buckets. In practice, to fill 4^*m*^ buckets, we compute larger minimizers of length *m*+*s* and hash them with a function *hash shuffle* that is distinct from the hash function used to order minimizers. Each (*m*+*s*)-minimizer joins one of the 4^*m*^ buckets according to the formula *hash shuffle*(*large minimizer*)%4^*m*^. This way we associate to each *k*-mer one of the 4^*m*^ buckets relying on its (*m*+*s*)-minimizer instead of its *m*-minimizer. Thus each bucket contains, on average, 4^*s*^ sub-buckets of (*m*+*s*)-minimizer, granting an improved balance of buckets.

#### 2.3.5 Sparse index

We can index super-*k*-mers approximate rank via sub-sampling to reduce the position encoding’s memory impact. The trade-off involves a smaller index for higher query time. More precisely, in order to save *b* bits per *k*-mer when encoding the rank, we indexes 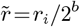 for each *k*-mer *k*_*i*_ of rank *r*_*i*_. From a given approximate rank, we have to check at most to 2^*b*^ super-*k*-mers during the query to ensure the queried *k*-mer exists. While scanning the *k*-mers of the different possible super-*k*-mers, we use the rank structure to check only indexed *k*-mers and avoid artefactual *k*-mers at the junction of two super-*k*-mers. In practice, to search the super-*k*-mers of rank *r*_*i*_ and *r*_*i*+1_, we effectively check the *k*-mers in the range [*position*(*r*_*i*_)..*position*(*r*_*i*+1_) − *k*] for the first super-*k*-mer and the range [*position*(*r*_*i*+1_)..*position*(*r*_*i*+2_) −*k*] for the second one. We experimentally show (Table 2) that up to *b* =4, the additional time has a limited impact on the query throughput.

**Table 2:**
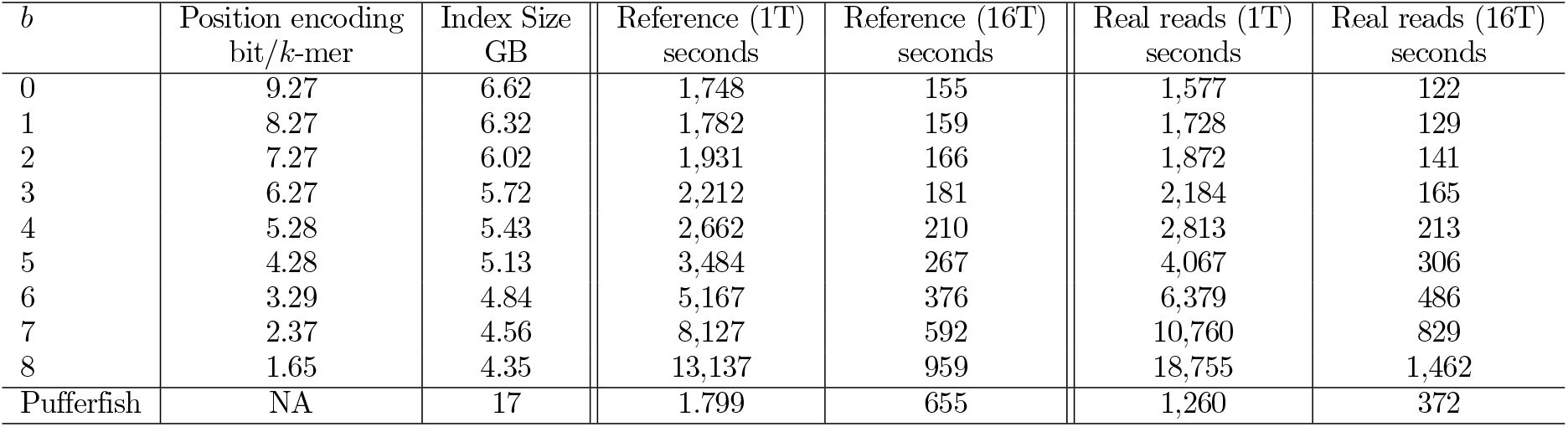
Influence of the sub-sampling parameter on the query time. We report the amount of memory used by the positions divided by the number of *k*-mers, the time required to query the real dataset SRR5833294 with one thread, the time required to query the real dataset with 12 threads, the time required to query all the *k*-mers of the reference graph and the time required to query this graph using 12 threads. Pufferfish queries cannot benefit from multiple threads.

We note that Pufferfish also implements a sparse index with a different sub-sampling strategy. This strategy associates positions to a subset of *k*-mers and how to find the nearest indexed *k*-mer otherwise. We could implement this strategy in the BLight index, but we favor our scheme for simplicity and efficiency with low sub-sampling ratios.

## 3 Results

We benchmark our structure on three different use cases. First, we select large, complex, reference genomes of increasing sizes (starting from the human that represents a moderate reference size, up to the currently largest available at NCBI, i.e., the axolotl *Ambystoma mexicanum*, with 32 Giga base pairs) in order to demonstrate how the structure scales to these objects. We use these datasets to demonstrate how the minimizer size impacts our structure’s performances and illustrate the trade-off obtained when using our index’s sparse version. In this application, we compare the current state-of-the-art approach designed to index such data, Pufferfish. We also show how our index can also benefit from other SPSS than the unitigs of the compacted de Bruijn graph. We compared the index performances on raw de Bruijn graph and using UST [21] showing a significant gain in performance using this approach.

Second, we demonstrate how our method can handle the indexation of raw reads datasets and show an example of potential applications of such an index. We show that BLight can index massive *k*-mer sets from raw NGS datasets, either from metagenomics or transcriptomic samples, up to an enormous soil sample containing more than 19 billion distinct *k*-mers.

We also used BLight to associate to each *k*-mer its number of occurrences across the datasets to show a library’s straightforward application. To do so, we selected a dataset from TARA Oceans samples [29], and compare BLight to two lightweight recent *k*-mer abundance indexes: Short Read Connector (SRC) [18] and Squeakr [30].

All experiments were performed on a single cluster node running with Intel(R) Xeon(R) CPU E5-2420 @ 1.90GHz with 192GB of RAM and Ubuntu 16.04.

### 3.1 Indexing up to top-largest reference genomes

#### Selected genomes

To assess the impact of the proposed minimizer partitioning, we built a de Bruijn graph (*k* =31) from several reference genomes and built the BLight index with different minimizer sizes on their graphs:

- The human reference genome (GRCh38.p12) of 3.2 Giga base pairs, counting 2.5 billion distinct *k*-mers and constructed with Bcalm2 [20] using 12 CPU hours and 6.6GB of RAM.
- The latest assembly of *Pinus taeda* (GCA 000404065.3) of 22 Giga base pairs, counting 10.5 billion distinct *k*-mers and constructed with Bcalm2 using 68 CPU hours and 17.3GB of RAM.
- The latest assembly of *Ambystoma mexicanum* (GCA 002915635.2) of 32 GbpGiga base pairs counting 18.3 billion distinct *k*-mers and constructed with Bcalm2 using 107 CPU hours and 44.1GB of RAM. To compare BLight to Pufferfish, we also included the bacterial genomes graph from Pufferfish paper, constructed from more than 8000 bacterial genomes counting 5.4 billion *k*-mers.

#### Index construction

We constructed the BLight index on the previously mentioned genomes graphs with several minimizer sizes to assess the minimizer size’s impact. We report in Figure 4 the memory necessary to encode the graph super-*k*-mers, to encode the *k*-mers positions, the total amount of memory taken by the index, and the actual maximal memory peak during construction. All values are in bits per *k*-mer.

**Figure 4.**
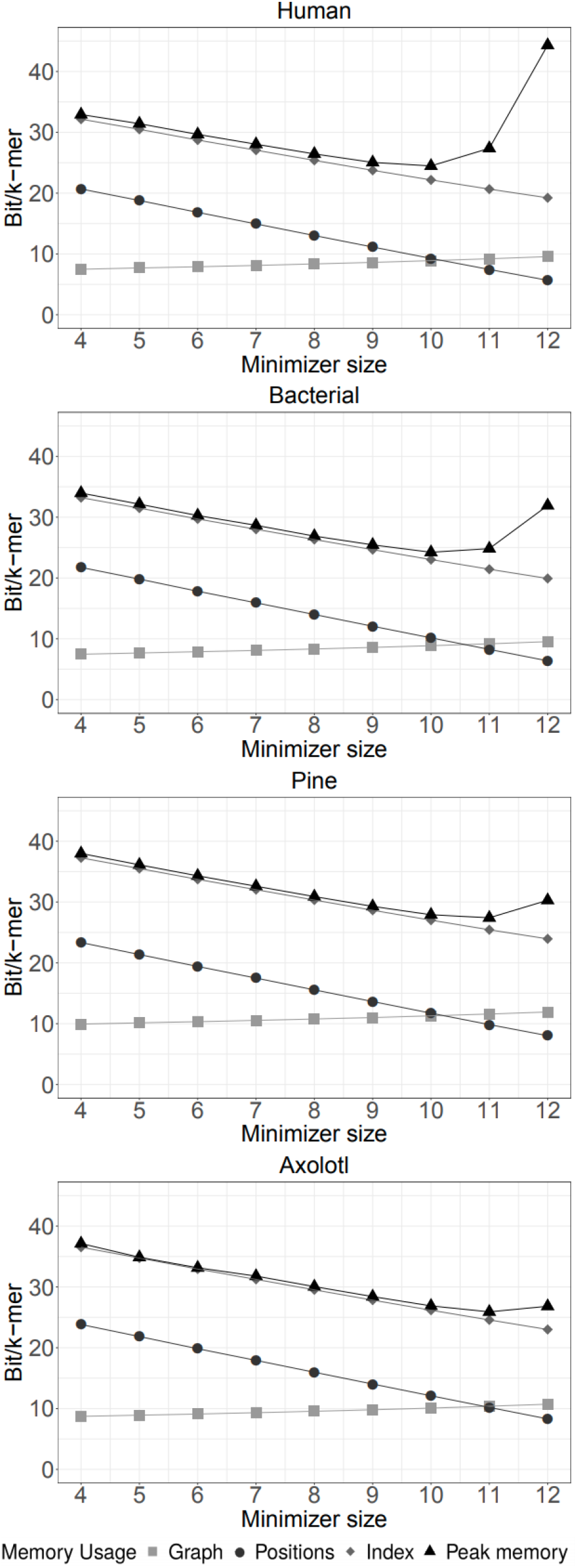
Detailed memory usage in bits per *k*-mer respectively on the human, bacterial genomes, pine and axolotl graphs during the BLight index construction.

We globally observe that a larger minimizer size leads to a slight augmentation of the memory needed to encode the graph sequences and decrease the memory required for position encoding. With uniform bucket filling, we expect the bits required to encode a *k*-mer position to be 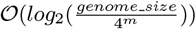 resulting in an expected gain of 2 bits per *k*-mer for increasing the minimizer size of one. Interestingly, the observed results are close to this trend in practice on our indexed genomes. We observe that larger minimizers lead globally to smaller indexes because of the reduction of the positions encoding. With a large minimizer size, we can reach the point where the graph itself is more memory expensive than the positions.

However, we also observe the influence of the exponential overhead that becomes prohibitively high after 12. We also notice that large minimizer sizes suit massive sets better as the high overhead may be too expensive to reduce memory usage. In practice, the minimizer size that achieves lowest memory usage is 10 for the human but 11 for the Axolotl.

We summarized the best results obtained with BLight compared to Pufferfish in Table 1. We built Pufferfish and Blight indexes directly on the compacted de Bruijn graphs to ensure a fair comparison. Pufferfish performances do not include the graph construction step and are using the fastest dense mode. We show that BLight constructs its index using substantially less memory than Pufferfish construction steps and the Pufferfish index itself. We ran all experiments using 12 threads to raise the construction speed. For comparison, the mono-thread construction of the Blight human index lasted 60 minutes. We also show improved construction time, and we want to highlight that, unlike Pufferfish, no pre-processing is needed on the graph for the index construction since we only require the unitig sequences in a FASTA file.

**Table 1:** Comparison of the maximal memory usage in GB and the Wall-clock time in minutes used during construction with BLight and Pufferfish from the de Bruijn graph. We were unable to construct Pufferfish’s index for the pine and the axolotl genomes due to memory limitation.

| Genome | Time (min) | Index Size (GB) | Memory usage (GB) | Bits/$k$-mer |
| --- | --- | --- | --- | --- |
| Pufferfish Human | 69 | 17 | 27.4 | 55.59 |
| BLight Human | 10 | 6.62 | 7.4 | 22.3 |
| Pufferfish Bacterial genomes | 187 | 40 | 75 | 114.8 |
| BLight Bacterial genomes | 21 | 15.73 | 16 | 24.23 |
| BLight Pine | 39 | 33.84 | 35.7 | 27.41 |
| BLight Axolotl | 76 | 53.62 | 62.4 | 26.88 |

However, the comparison is not fair as Pufferfish was designed to be a reference index to associate to each *k*-mer its positions in a reference genome. Therefore Pufferfish needs to keep this information during the construction of the graph and in the index. This information’s memory surplus is hard to estimate, but we can observe our partitioning method’s interest in the used memory, as a small minimizer size is similar to a strategy without partition.

#### Query time and impact of sub-sampling factor

We compared our throughput with Pufferfish’s (using the fastest “dense” mode) in Table 2. We observe that our query is similar to Pufferfish’s on a single CPU. However, using multiple threads significantly raise Blight throughput, outperforming Pufferfish. Either way, both indexes propose a very high throughput of the order of the million queries per second. BLight reaches 1.4 million *k*-mer queries per second on a single core and 16.1 million *k*-mer queries per second using 12 cores. Using a sub-sampling factor can mitigate memory usage at the expense of the query performances. However, we observe that below *b* =4, the impact on the query time is low.

#### BLight as a de Bruijn graph representation

BLight can be used as a high-throughput de Bruijn graph representation but is not competitive compared to space-efficient graph structures. We computed the theoretical bound for each species indicated by Conway & Bromage (CB) [31] for *k* =31 in bits per *k*-mer: human 32.2, pine 30.15, axolotl 29.35 and bacterial genomes set 31.11. Contrary to our representation and Pufferfish’s, CB does not consider the possibility to combine the information of multiple adjacent *k*-mer. This fact explains why we can use less space than CB. However, our structure and Pufferfish’s use 𝒪 (*log*(*genome size*)) bits per *k*-mer, thus could be higher than CB on larger genomes. Other FM-index based methods such as DBGFM maintain compressed representations of the de Bruijn graph, thus can be more space-saving than ours. The proposed methods provide a different trade-off as we expect a faster query time in practice.

#### Benefit of Spectrum-preserving string sets

In BLight, our first choice was to represent *k*-mer sets in an exact and partitioned way using super-*k*-mers from unitigs. However, BLight can construct its index from any Spectrum-preserving string sets without duplicate. We recall that in this work, we are interested in SPSS that represents a set of *k*-mers and will refer to them, and will not take into account multi-sets. Unitigs are one SPSS, super-*k*-mers of unitigs are another [14]. Two other equivalent SPSSs schemes, UST [21] and simplitigs [32], longer than unitigs, were recently independently proposed. The motivation of those two developments is that unitigs are not the most concise representation of a set of *k*-mer as we can further compact the compacted de Bruijn graph without altering the *k*-mer set they represent.

We used UST to further compact the human and pine unitigs of Table 1 and compared the amount of memory used on the unitigs versus UST SPSS and report the result in Table 3. We observe improvements in the memory required to encode both the graph sequences and the *k*-mer positions resulting in significantly smaller indexes (from 30 to 26 GB on the Pine genome with a minimizer size of 12). This table shows the benefit of using SPSS to represent and index *k*-mer set more concisely and how BLight can benefit from those approaches.

**Table 3:** Space usage of BLight indexes on raw unitigs compared to unitigs compacted with UST.

| File | Minimizer<br>size | Graph<br>bits per $k$-mer | Position<br>bits per $k$-mer | Total<br>bits per $k$-mer |
| --- | --- | --- | --- | --- |
| Human | 8 | 8.1 | 13.9 | 26 |
| Human UST | 8 | 7.8 | 12.9 | 24.7 |
| Human | 10 | 8.6 | 10.1 | 22.7 |
| Human UST | 10 | 8.3 | 9.1 | 21.5 |
| Pine | 8 | 10.5 | 16.4 | 31 |
| Pine UST | 8 | 8.6 | 15.2 | 27.8 |
| Pine | 10 | 11 | 12.6 | 27.6 |
| Pine UST | 10 | 9.2 | 11.4 | 24.5 |
| Pine | 12 | 11.6 | 8.9 | 24.5 |
| Pine UST | 12 | 9.8 | 7.7 | 21.5 |

### 3.2 Indexing *k*-mer from raw reads

#### Indexing metagenomic datasets

In this section, we assess the ability of the BLight structure to index complex metagenomic datasets. The following experiment is contrasting with the bacterial genome collection used in Figure 4. In the previous experiment, the de Bruijn graph is constructed from reference genomes, while here we use raw datasets, yielding very different graph topologies. They are highly fragmented because of missing regions, sequencing errors, and related genomes present [33]. We first chose to index all non-unique *k*-mers of a TARA sample^2^ containing five datasets ERR1712199, ERR1711907, ERR599280, ERR562434, and ERR1718455. BLight was able to build an index from the graph of 7,540,111,917 distinct *k*-mers constructed from Bcalm2. The construction from unitigs lasted less than an hour and used 24.7 GB, representing 27.1 bits per *k*-mer. This memory usage is higher than the bacterial genomes despite having a comparable amount of *k*-mers. This surplus is due to the expected graph fragmentation of a raw metagenomic sample that raises the nucleotide amount needed to encode the graph (see Minimizer size and fragmentation section).

We also indexed all non-unique *k*-mers of a massive soil sequencing from DOE/JGI Great Prairie Soil Metagenome Grand Challenge^3^ counting 19,289,529,788 distinct *k*-mers. The construction lasted less than a day and used 63.8GB, representing 26.5 bits per *k*-mer.

##### Indexing transcriptomic datasets

In this section, we assess the ability of the BLight structure to index complex transcriptomic datasets. As most transcriptomic sequencing experiments contain a “low” amount of distinct *k*-mers we choose to index all distinct *k*-mers from the 2,585 datasets used for the Sequence Bloom Tree benchmark [10] using their specific filtering method. We build the union de Bruijn graph from all the files using Bcalm2 with *k* = 31 and *k* = 21. The graphs have 4,425,877,751 distinct 31-mers and 3,903,590,501 distinct 21-mers respectively. Both indexes were built within 30 minutes and used 16.83 GB (30.4 bits per 31-mers) and 16.31 GB (33.4 bits per 21-mers), respectively.

#### 3.3 Application example: storing *k*-mer counts

In this experiment, we provide a proof of concept index to associate to each *k*-mer its abundance within a dataset. We compare a simple usage of BLight through a *k*-mer abundance index snippet, with two methods from state-of-the-art that allow large scale *k*-mer to abundance association.

Squeakr [30] is a *k*-mer counter based on a quotienting hashing technique, and Short Read Connector counter (SRC) [18] is based on an MPHF as BLight. The proposed snippet constructs the compacted de Bruijn graph of the input file with Bcalm2, constructs a BLight index from it, allocates a byte for each distinct *k*-mer and then queries all *k*-mers of the input file to increment its count. We report the performances of the three tools on a large marine metagenomic dataset from TARA used the previous metagenomic experiment (ERR599280), counting 37 billion bases and 189 million reads. All tools are used in exact mode and presented in Table 4. Since this dataset counts only 329,265 *k*-mers with a count above 255 and barely 325 *k*-mers with a count above 65,536 we choose to limit SRC and BLight maximal count to 255 for a cost of one byte per *k*-mer, while Squeakr handles very larger counts. Handling abundant *k*-mers (above 255) would require an overhead of one byte per *k*-mer and handling very abundant *k*-mers (above 65,536) would require an overhead of three bytes per *k*-mer. If the memory usage of BLight is significantly reduced compared to the other methods, the index construction is way slower than SRC or Squeakr. However, here Blight performs a query for each *k*-mer of the dataset to count its abundance, representing an excessive construction time. A real implementation should rely on an efficient *k*-mer counter and parse a *k*-mer counting result to initialize the *k*-mer abundance as SRC. Moreover, SRC or Squeakr inexact modes are expected to be more space-efficient but will yield false positives and inexact results.

**Table 4:** Performance comparison of an abundance index construction on the ERR599280 metagenomic dataset.

| Tool | Peak memory (GB) | Time (hh:mm) |
| --- | --- | --- |
| Squeakr exact | 185.4 | 15:11 |
| SRC counter exact | 44.20 | **05:12** |
| BLight count | **20.8** | 19:49 |

## 4 Conclusion and future work

In this work, we propose BLight, a low-cost, high-throughput, and exact associative structure for indexing *k*-mers, relying on de compacted Bruijn graphs or other efficient SPSS. Based on efficient hashing techniques and light memory structure, we argue that the proposed index has a very interesting time/memory compromise, performing millions of queries per second, and using less than 32 bits per *k*-mer on our large-scale experiments. In comparison, we show that the closest method, Pufferfish, reaches a different compromise. While not benefiting from the gain in space due to partitioning, it optimizes the storage of the *k*-mers positions within a reference genome, making Pufferfish more suitable for applications such as alignment on a reference. Conversely, we designed BLight with a more generic application range in mind. We showed that BLight can index the largest available genomes to date using a reasonable amount of memory while outperforming state of the art methods in both construction time and memory. We also showed it could be relevant and efficient on raw transcriptomic and metagenomic sequencing data.

Being able to work on massive datasets is often a challenge in sequence analysis. By providing an efficient structure able to run on medium clusters or laptops, we hope to democratize and lower this type of analysis’s inherent cost. We thus believe that a vast number of methods could rely on and benefit from this structure due to its broad application spectrum. To that extent, we implemented a user-friendly library and different snippets to allow our method to be usable in practical cases. The challenge of indexing colored de Bruijn graphs [34] (or more generally to answer large sequence search problems as defined in [10]) have caught the interest of a community and could be a direct application of this work. For example, BLight is successfully integrated as an indexing structure in REINDEER [14], a *k*-mer data structure that enables the quantification of query sequences in thousands of raw read samples.

The main caveat of the proposed structure is that the index cannot include new sequences after its construction. The static aspect allows the index to be extremely memory-efficient, which would be hard to achieve with a dynamic structure. A possible continuation of this work would be a dynamic structure that follows the main idea of BLight, using multiple dynamic indexes partitioned by minimizers. This partitioning should improve the data locality and, therefore, the performance of such a structure. Another way to pursue this work would be to propose a pseudo-dynamic structure using a static and a dynamic structure for the last elements inserted. At some point, in a way to guarantee a constant average insertion time, a complete rebuild could be performed to construct a static index on all elements. More generally, we could bring a vast amount of improvements to the proposed backbone structure. We could design specific minimizer schemes to obtain the smallest possible buckets, supplementary structures to sort and rank the super-*k*-mers could allow faster query or reduced position encoding. Such optimizations could improve the global resource usage of the index or provide different time/memory trade-offs.

## Acknowledgements

We want to thank Rob Patro, Fatemeh Almodaresi, Tatiana Rocher, and Rayan Chikhi for their support and exciting discussions on this project. The ANR Transipedia supported this work (ANR-18-CE45-0020).

## Footnotes

1 github.com/tlk00/BitMagic

2 https://www.ebi.ac.uk/ena/data/view/SAMEA2620556

3 www.mg-rast.org/mgmain.html?mgpage=project&project=mgp6377

